# Ageing impairs the airway epithelium defence response to SARS-CoV-2

**DOI:** 10.1101/2021.04.05.437453

**Authors:** Alexander Capraro, Sharon L Wong, Anurag Adhikari, Katelin M Allan, Hardip R Patel, Ling Zhong, Mark Raftery, Adam Jaffe, Malinna Yeang, Anupriya Aggarwal, Lindsay Wu, Elvis Pandzic, Renee M Whan, Stuart Turville, Orazio Vittorio, Rowena A Bull, Nadeem Kaakoush, William D Rawlinson, Nicodemus Tedla, Fatemeh Vafaee, Shafagh A Waters

## Abstract

Age-dependent differences in the clinical response to SARS-CoV-2 infection is well-documented^1–3^ however the underlying molecular mechanisms involved are poorly understood. We infected fully differentiated human nasal epithelium cultures derived from healthy children (1-12 years old), young adults (26-34 years old) and older adults (56-62 years old) with SARS-COV-2 to identify age-related cell-intrinsic differences that may influence viral entry, replication and host defence response. We integrated imaging, transcriptomics, proteomics and biochemical assays revealing age-related changes in transcriptional regulation that impact viral replication, effectiveness of host responses and therapeutic drug targets. Viral load was lowest in infected older adult cultures despite the highest expression of SARS-CoV-2 entry and detection factors. We showed this was likely due to lower expression of hijacked host machinery essential for viral replication. Unlike the nasal epithelium of young adults and children, global host response and induction of the interferon signalling was profoundly impaired in older adults, which preferentially expressed proinflammatory cytokines mirroring the “cytokine storm” seen in severe COVID-19^4,5^. *In silico* screening of our virus-host-drug network identified drug classes with higher efficacy in older adults. Collectively, our data suggests that cellular alterations that occur during ageing impact the ability for the host nasal epithelium to respond to SARS-CoV-2 infection which could guide future therapeutic strategies.

## Introduction

Severe acute respiratory syndrome coronavirus 2 (SARS-CoV-2) is the causative agent of the COVID-19 pandemic^6^. Clinical presentation is highly heterogeneous, ranging from asymptomatic infection to critically ill patients with respiratory failure^7^. The single greatest risk factor for disease severity and mortality is age. Children experience relatively mild symptoms^1,2,8–10^, with a case fatality rate of 0.026% under the age of 9, compared to an increased risk of severe illness in older groups, with a case fatality rate of over 13% in those aged over 80 ^3^. This trend was also observed during the 2002-4 SARS epidemic^11^, and contrasts the 1918 H1N1 and 1968 H3N2 influenza epidemics, where mortality was concentrated in younger adults^8^. Despite this clear trend, the reasons for increased mortality from COVID-19 with ageing are uncertain.

The airway epithelium is the first site exposed to SARS-CoV-2 and is central to the host defence and innate immune response to infection. Ciliated and goblet cells are preferentially infected as they have the highest expression of the SARS-CoV-2 entry receptor angiotensin-converting enzyme 2 (ACE2)^12^. Upon entry, viral RNA normally triggers an innate immune response involving the transcription of antiviral and proinflammatory cytokines^13^ including type I and III interferons (IFNs) that induce the expression of IFN-stimulated genes (ISG) which restrict viral replication^14^. A key feature of coronaviruses is the ability to evade this innate immune response, primarily through the inhibition of IFN production and downstream signalling. This aberrant immune response is characterised by cytokine release syndrome (CRS) and dysregulated interferon response^4,5,15,16^, resulting in secondary bacterial pneumonia, acute respiratory distress syndrome (ARDS), multi-organ failure, and death^17^. Explanations for age-related differences in COVID-19 outcomes include differential expression of ACE2, altered responses from circulating immune cells, background co-morbidities and low-grade chronic inflammation (“inflammaging”). However, whether cell-intrinsic changes in the airway epithelium that occur with ageing contribute to these differences is unclear.

### Decreasing viral load in differentiated nasal epithelial cultures with ageing

To compare age-dependent mucosal innate immune responses to SARS-CoV-2 infection, we established well-differentiated human nasal epithelial cultures (hNEC) at air liquid interface (ALI) from nasal brushings of 13 participants, comprising six children (5.40±3.97 y/o), four young adults (31.89±4.14 y/o), and three older adults (58.14±3.30 y/o) (**Fig. 1A, Table S1**). Since SARS-CoV-2 displays a decreasing infectivity gradient from the upper to the lower respiratory tract^20^, hNECs from the child participants were also compared with participant-matched bronchial epithelial cells (hBEC). We confirmed the differentiated cultures retained the expected pseudostratified morphology of mucociliary epithelium (**Extended Data Fig. S1A)** and function. We observed a decline in ciliary beat frequency (CBF) in cultures from older adults that is consistent with known physiological changes with ageing^21,22^ (**Fig. 1B**). This provided functional evidence that the age-related properties of the hNEC participant was retained. In addition, both transcriptome and proteome mock-infected data showed significant age-dependent correlation with positive (adhesion and stress response) and negative (mitochondria and replication) age-associated genes reported for the ageing human lung^23^ (**Extended Data Fig. 1B-C**).

**Figure 1.**
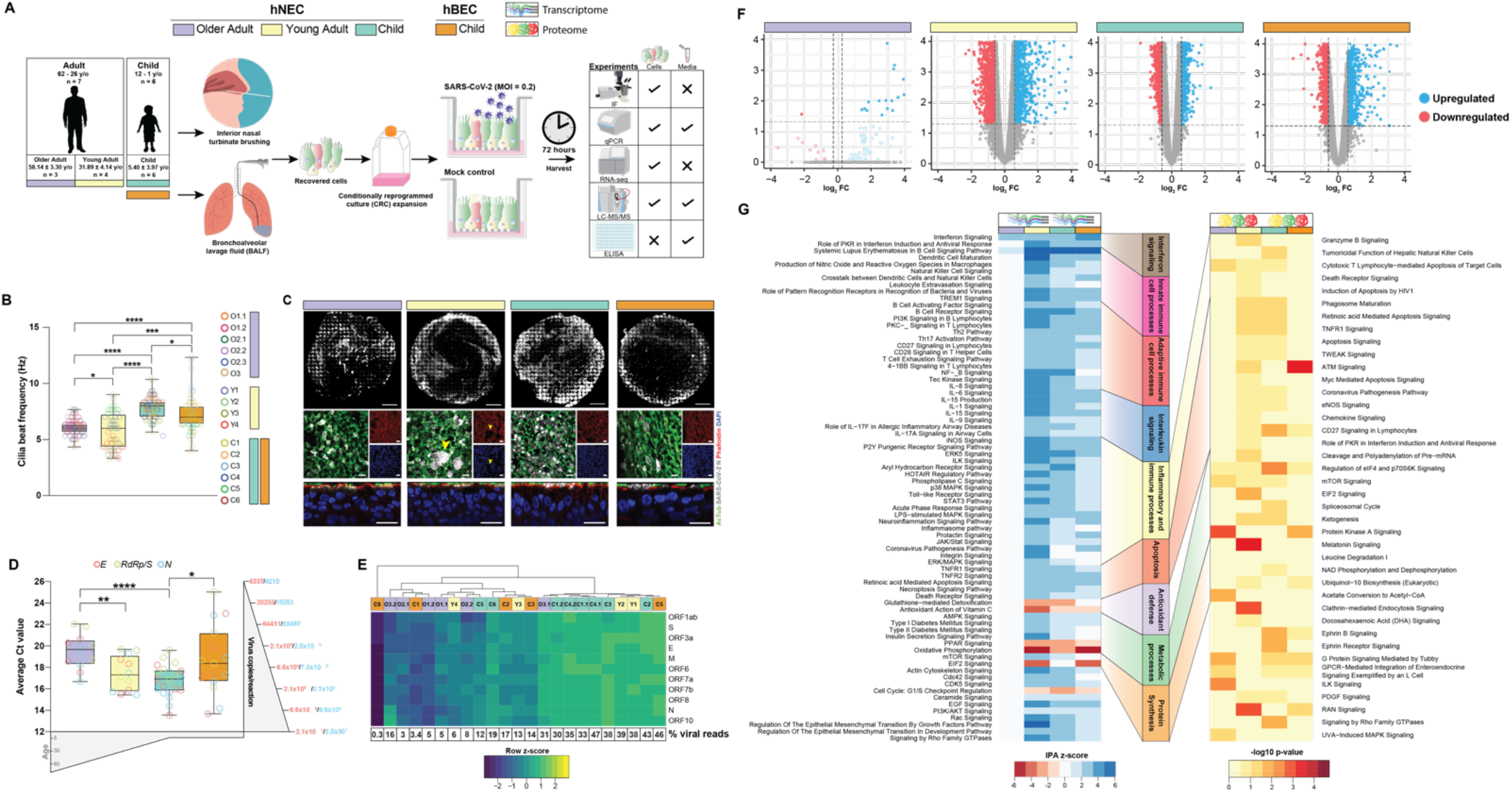
Differentiated human airway epithelial cultures have age-dependent viral load and transcriptional and proteomic response to SARS-CoV-2. **(A)** Schematic of study. Human nasal epithelial cells (hNEC) were harvested from the inferior nasal turbinate from adults (26 - 62 years old, n = 7) and children (1-12 years old, n = 6). Human bronchial epithelial cells (hBEC) were harvested only from matched child participants. Harvested cells were subject to conditional reprogramming culture (CRC) expansion. The reprogrammed cells were differentiated at air-liquid interface (ALI). Once matured (21 days), hNECs and hBECs were apically inoculated with PBS (mock) or SARS-CoV-2 (MOI: 0.2). Cultures were harvested and analysed 72 hours post-infection (hpi). **(B)** Mean cilia beat frequency (CBF) measurements (Hz) of the fully differentiated mature human airway epithelial cells (hAEC). Child, *n* = 5; young adult, *n* = 4; older adult, *n* = 6; paired two-tailed Student’s t-test. **(C)** Immunofluorescence (IF) staining of SARS-CoV-2 infected hNECs from representative older adults (donor O3), young adults (donor Y1) and children hNECs and hBECs (donor C2) 72 hpi. Top: Representative normalised 25×25 tilescan binary masked images of SARS-CoV-2 nucleocapsid positive clusters (grey) on the apical layer of the culture. 20x/0.5 objective. Scale bars = 1000μm. Middle: Top view; Bottom: Side-view of SARS-CoV-2 nucleocapsid (SARS-CoV-2+ cells, grey), acetylated tubulin (ciliated cell marker, green), phalloidin (actin, red) and DAPI (nucleus, blue). Yellow arrows indicate syncytia. 63x/1.4 oil immersion objective. Scale bars = 20μm. **(D)** Average Ct values of SARS-CoV-2 *E*, *RdRp*/*S* and *N* genes in SARS-CoV-2-infected cultures as measured with qPCR. Respective plasmid copies per reaction shown on the right y-axis for *E* and *N* gene. Child hNEC, *n* = 8; child hBEC, *n* = 5, young adult, *n* = 6; older adult, *n* = 6; unpaired two-tailed Student’s t-test. **(E)** Relative expression of all SARS-CoV-2 genes in SARS-CoV-2-infected hAECs in RNA-seq. Top row indicates different donors. Each tile shows the normalized expression of a SARS-CoV-2 gene (Z-score) ordered by genomic location. Colour key shows the relative expression in each sample (yellow, high; blue, low). **(F)** Volcano plots of differentially expressed genes between mock-infected and SARS-CoV-2-infected hAECs in children, young adults, older adults. Each dot indicates an individual gene with the colour indicating either upregulation (blue) or downregulation (red) relative to mock. Dotted lines indicate the significance cut-off (FDR < 0.05, fold change > |1.5|). **(G)** Top enriched canonical pathways of differentially expressed genes (left panel) and proteins (right panel) within each age group determined by IPA. Red-blue heatmap colour indicates the IPA z-score for each pathway within each age group, with positive (blue) z-scores indicating predicted activation and negative (red) z-scores indicating predicted inhibition. White-red heatmap colour indicates the −log_10_ p-value of each canonical pathway. Boxplots indicate the first and third quartiles with middle horizontal line indicating the median and external horizontal lines indicating minimum and maximum values. * *p*-value < 0.05, ** *p*-value < 0.01, *** *p*-value < 0.001, **** *p*-value < 0.0001.

Next, the apical surface of these cultures was infected with SARS-CoV-2 at 0.2 multiplicity of infection (MOI). Viral entry and internalisation was confirmed at 72 hours post-infection (hpi) using immunofluorescence (IF) staining of the viral nucleocapsid protein (SARS-CoV-2 N). The protein was present within the cytoplasmic compartment and manifested in clusters which were limited to the apical top layer of cultures in all age groups (**Fig. 1C)**. Consistent with the decreasing infectivity gradient down the respiratory tract^20^, approximately 20% of the top layer of cells were positively stained for SARS-CoV-2_N in hNECs created from the adult and child participants, whereas only 8% of the top layer of child hBECs were positive (**Fig. 1C, Table S3**).

Viral load was assessed from infected cells using quantitative PCR (qPCR) targeting viral cDNA, RNA sequencing (RNA-seq) and protein quantification by LC/MS-MS. Surprisingly, viral transcript levels assessed by qPCR were significantly lower (*p* < 0.01) in hNECs from older adults than those of children and young adults (**Fig. 1D**). This result was confirmed with RNA-seq with ~9% of reads from SARS-CoV-2 in hNECs from older adults compared to ~28% in children and young adults (**Fig. 1E, Extended Data Fig. 1D-E**), and by the differential abundance of SARS-CoV-2 proteins detected by unbiased proteomics (**Extended Data Fig. 1F**). As with the results of viral staining and the gradient of infection observed clinically, SARS-CoV-2 transcription in hBECs were significantly lower than those in patient-matched hNECs despite infection at the same MOI (**Fig. 1C-E**). This demonstrates that ageing influences SARS-CoV-2 viral load 72 hpi in the hNEC model (**Extended Results**).

### Transcriptional responses to SARS-CoV-2 infection are severely blunted in older adults

Given the age-dependent variation in viral load, we next surveyed global transcriptomic and proteomic responses to infection. A strong age-dependent response was evident when host transcriptional and proteomic profiles in SARS-CoV-2 infected cells were compared with their respective uninfected controls (**Fig. 1F**, **Extended Data Fig. 2A-C**, **Table S4, S5**). Children and young adults hNECs displayed robust, global proteomic and transcriptional responses to SARS-CoV-2 infection, with 3,240 (20% of all expressed genes) and 2,466 (15%) differentially expressed genes (DEG), respectively. In contrast, the transcriptional response to infection in older adult hNECs was almost completely absent, with only 24 (0.015%) DEGs (**Fig. 1F**). Even with a less stringent cut-off (*p*<0.05 rather than FDR<0.05), only 422 (2.6%) DEGs were identified (**Fig. 1F**) This was unlikely due to the low viral load in the older adult hNECs as child hBECs with a similarly low viral load (~9% and ~12%, respectively; **Fig. 1C-E**) showed a robust transcriptional response to SARS-CoV-2 infection (DEGs=1,797, 11%; **Fig. 1F, Extended Data Fig. 2A, Table S4**). The proteomes of these samples followed a similar trend (**Extended Data Fig. 2B-C, Table S5**) and was highly correlated with transcript levels in both mock and SARS-CoV-2 infected cultures (Spearman’s *ρ*=0.483 and 0.486, *p*<0.05, respectively; **Extended Data Fig. 2D**).

**Figure 2.**
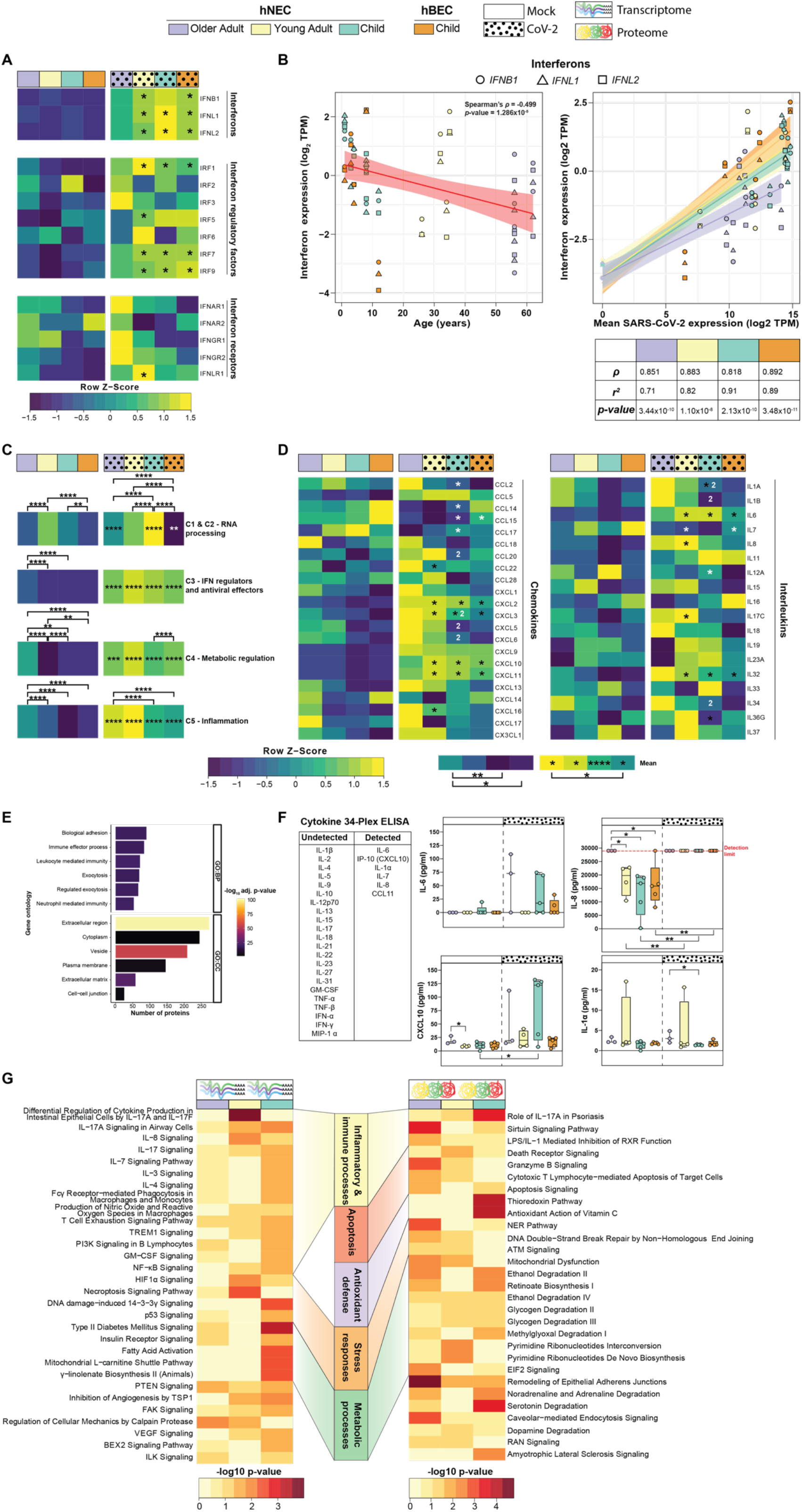
Interferons and cytokines are induced in an age-dependent manner to SARS-CoV-2 infection. **(A)** Heatmap of the relative gene expression of interferons, interferon regulatory factors and interferon receptors. **(B)** Spearman’s correlation of the gene expression of type I (*IFNB1*) and III *(IFNL1* and *IFNL2*) interferons (log_2_ transcripts per million (TPM)) and SARS-CoV-2 gene expression (log2 TPM) expression in all cultures (bottom panel) and age in SARS-CoV-2-infected cultures (top panel). Shapes show the three interferon genes with black bordered shapes indicating SARS-CoV-2 infected cultures. Heatmap of the **(C)** relative mean expression of Interferon-Stimulated Genes (ISG) within functional clusters as defined by Mustafavi et al (2016), and **(D)** relative expression of cytokines (chemokines and interleukins). For heatmaps colour key shows the relative expression (row Z-score; yellow, high; blue, low). * significantly upregulated (black) and downregulated (white) in SARS-CoV-2 infection relative to mock. (1) significantly upregulated and (2) downregulated relative to the two other age groups within the same condition. Mock, child hNEC, *n* = 5; child hBEC, *n* = 5, young adult, *n* = 4; older adult, *n* = 5; virus, child hNEC, *n* = 8; child hBEC, *n =* 5, young adult, *n* = 4; older adult, *n* = 6; two-tailed Wilcoxon rank sum test with Bonferroni correction. * adjusted *p*-value < 0.05, ** < 0.01, *** < 0.001, **** < 0.0001. **(E)** Enriched biological process (BP) and cellular compartment (CC) gene ontologies (GO) of identified proteins in the basolateral media of all cultures. Colour indicates the −log_10_ p-value for each pathway. **(F)** Cytokine and Chemokine 34-plex ELISA of the basolateral media of mock and SARS-CoV-2 infected cultures. Red dotted line indicates the detection limit of the assay. Child hNEC, *n* = 5; child hBEC, *n* = 5, young adult, *n* = 4; older adult, *n* = 4; unpaired two-tailed Student’s t-test. Boxes indicate the first and third quartiles with middle horizontal line indicating the median and external horizontal lines indicating minimum and maximum values. * *p*-value < 0.05, ** < 0.01. **(G)** Canonical pathway analysis of age-associated genes identified in the transcriptome and proteome as determined by IPA. Colour indicates the −log_10_ p-value of each canonical pathway.

Pathway enrichment analysis revealed similar responses in hNECs and hBECs from children and young adults hNECs, with upregulation of interferon signalling, innate and adaptive immune response, apoptosis, cytokine, and inflammatory response pathways, while antioxidant defence, oxidative phosphorylation, and fatty acid metabolism were reduced (**Fig. 1G, Table S6, S7**). In older adults, only interferon signalling, and B cell signalling were altered (**Fig. 1G**), however, the less stringent cut-off (**Fig. 1F**) expanded this to include hypercytokinemia (cytokine storm), coronavirus pathogenesis and viral recognition pathways (**Extended Data Fig. 2E**). Notably, this trend was not restricted to coding transcripts, with differential expression of long non-coding RNAs (lncRNA) in young adult (127), child hNECs (164) and hBECs (92) compared to only a single lncRNA (*RN7SL1,* RNA component of signal recognition particle) differentially expressed in older adults (**Extended Data Fig. 2F, Table S8**; **Extended Results)**. Pathway analysis of proteomics data revealed similar pathways, however, lower numbers of differentially expressed proteins (DEP) restricted interpretation (**Fig. 1G, Table S7**).

### Dysregulated IFN signalling and antiviral responses to SARS-CoV-2 with ageing

A key feature of coronaviruses is the ability to evade antiviral IFN signalling^16,24^, with susceptibility to exogenous IFN treatment^16,25^ and the host ability to mount an IFN response important determinants of patient survival^26^. In line with this, type I (*IFNB1*) and type III (*IFNL1*, *IFNL2*) IFNs were induced by viral infection in cultures from children and young adults, but not in hNECs from older adults, with a similar trend observed for IFN regulatory factors (IRFs) (**Fig. 2A**). In accordance with this, IFN expression in SARS-CoV-2 infected cultures was moderately negatively correlated with age (Spearman’s *ρ*=-0.499, *p*<0.05; **Fig. 2B**). While IFN gene expression was strongly positively correlated with SARS-CoV-2 transcript expression across all age groups, the coefficient of determination was 20% lower in older adults compared to children and young adults suggesting that ageing leads to less effective induction of IFNs by SARS-CoV-2 (**Fig. 2B**).

IFN induction was associated with the expression of several hundred IFN-stimulated genes (ISG; **Fig. 2C, Extended Data Fig. 3A-B**). ISGs were grouped based on five functional clusters as defined by Mostafavi^27^; RNA processing (C1 and C2), interferon regulation and antiviral function (C3), metabolic regulation (C4) and inflammation (C5). Cluster C1 and C2 was profoundly lower in older adults compared to other age groups (adjusted *p*-value<0.0001), which follows the common RNA virus strategy to inhibit host RNA processing to restrict the host antiviral response^28^. Despite an impaired IFN response in older adults, baseline expression of C3, C4 and C5 clusters were significantly higher in uninfected older adults (adjusted *p*-value < 0.0001), and all were increased upon infection when compared to their mock counterparts (adjusted *p*-value < 0.05). In contrast, the mean expression of genes in the inflammation cluster was significantly higher in older adult than both child cultures (adjusted *p*-value < 0.0001). Notably, this exaggerated response in older adult was elicited by a subset of only 14/552 ISGs (**Extended Data Fig. 3A**), compared to the broad upregulation of ISGs in young adult (156/552) and child hNEC (132/552) and hBECs (109/552) with a similar trend in the ISG protein levels (**Extended Data Fig. 3B, Extended Results**).

**Figure 3.**
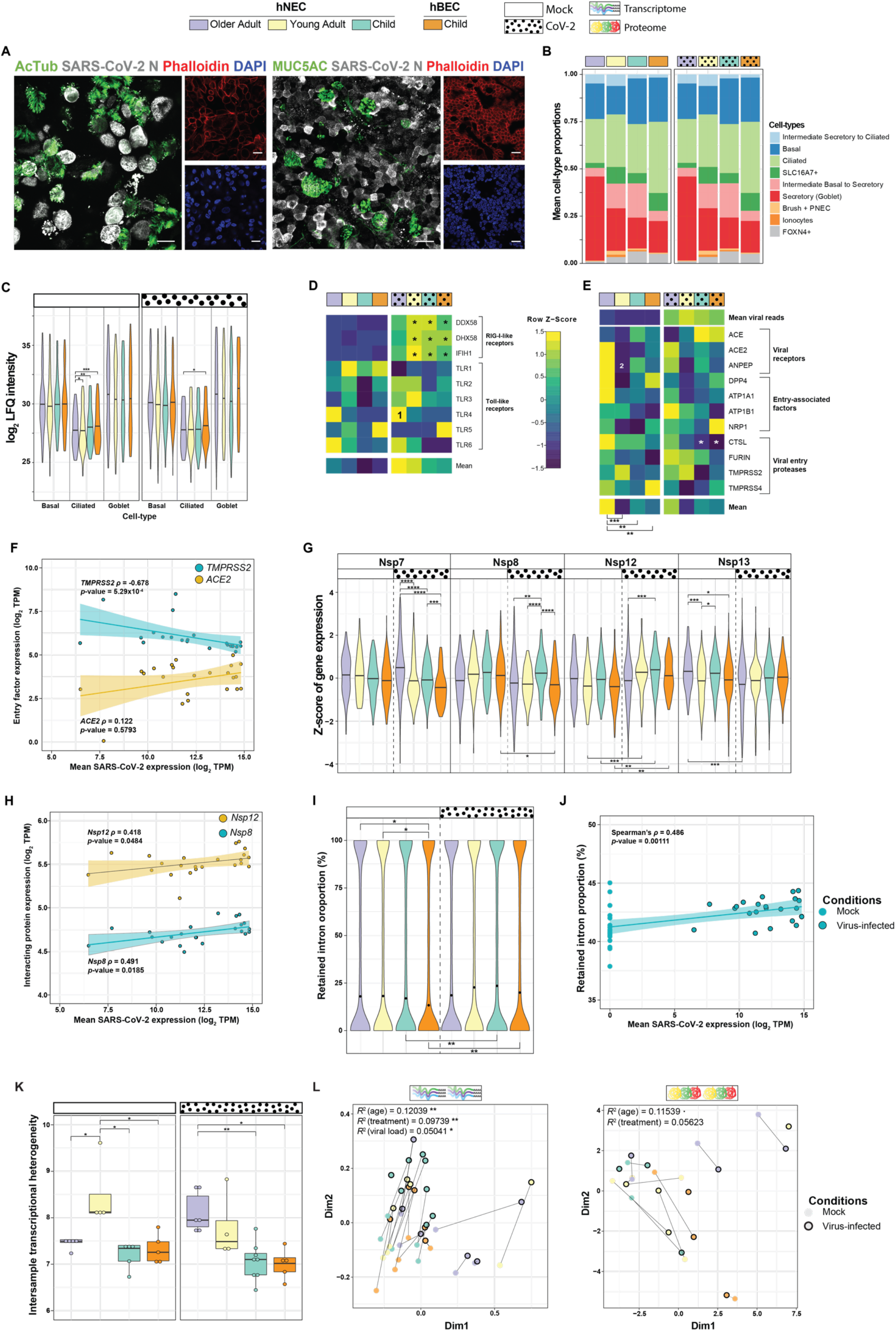
Host-virus interacting factors affect viral entry and replication. **(A)** Representative immunofluorescence images of differentiated SARS-CoV-2+ hNECs co-stained for acetylated tubulin (ciliated cells, green), MUC5AC (mucus-producing goblet cells, green), phalloidin (actin, red) and DAPI (nucleus, blue). 63x/1.4 oil immersion objective. Scale bars: 20μm. **(B)** Estimated cell-type composition of mock-infected and SARS-CoV-2-infected hAECs predicted by CIBERSORTx. Plot shows the mean cell-type proportions of all replicates within each culture with colours indicating cell-type. **(C)** Average log_2_ label-free quantification (LFQ) intensity of all identified markers for the major airway epithelial cell-types in mock-infected and SARS-CoV-2-infected hAECs. Mock, child hNEC, *n* = 2; child hBEC, *n* = 2, young adult, *n* = 4; older adult, *n* = 3; virus, child hNEC, *n* = 3; child hBEC, *n =* 3, young adult, *n* = 4; older adult, *n* = 4; two-tailed Wilcoxon rank sum test with Bonferroni correction. Relative mean expression of SARS-CoV-2 **(D)** pattern recognition and (**E**) entry factors in mock and SARS-CoV-2-infected hAECs. Each tile shows the normalized expression of a gene (Z-score). Mean expression were calculated across each age group and condition. Colour key shows the relative expression (yellow, high; blue, low). **(F)** Spearman’s correlation of mean *ACE2* and *TMPRSS2* gene expression (log_2_ transcripts per million (TPM)) and SARS-CoV-2 gene expression (log2 TPM) in SARS-CoV-2-infected hAECs. (**G**) Relative mean gene expression of host proteins that interact with the SARS-CoV-2 replication complex (Nsp7, Nsp8, Nsp12 and Nsp13). **(H)**Spearman’s correlation of mean gene expression of host proteins that interact with *NSP12* and *NSP8* (log_2_ TPM) and SARS-CoV-2 gene expression (log_2_ TPM) in SARS-CoV-2-infected cultures. **(I)**Retained intron proportion (%) for all genes with intron retention events across all age groups and conditions. Black dots indicate median for each group. **(J)** Spearman’s correlation of mean intron retention proportion (%) and SARS-CoV-2 gene expression (log_2_ TPM) in SARS-CoV-2-infected cultures. **(K)** Transcriptional noise of mock and SARS-CoV-2-infected cultures. Transcriptional noise was measured as the Euclidean distance between each sample and the mean within each age group for the 10% of the genes with the lowest coefficient of variation. Boxes indicate the first and third quartiles with middle horizontal line indicating the median and external horizontal lines indicating minimum and maximum values. **(L)** Principal coordinate analysis (PCoA) of the transcriptome (left) and proteome (right) in mock-infected and SARS-CoV-2-infected hAECs. Color indicates the age group and outline indicates condition. Statistical significance of the PCoA was performed with the *adonis* function within the *vegan* R package. Mock, child hNEC, *n* = 5; child hBEC, *n* = 5, young adult, *n* = 4; older adult, *n* = 5; virus, child hNEC, *n* = 8; child hBEC, *n =* 5, young adult, *n* = 4; older adult, *n* = 6; two-tailed Wilcoxon rank sum test with Bonferroni correction. For panels C, G, I and K: * adjusted *p*-value < 0.05, ** adjusted *p*-value < 0.01, *** adjusted *p*-value < 0.001, **** adjusted *p*-value < 0.0001. For heatmaps: * significantly upregulated (black) and downregulated (white) in SARS-CoV-2 infection relative to mock. (1) significantly upregulated and (2) downregulated relative to the two other age groups within the same condition.

### Inflammaging and heightened inflammatory response to SARS-CoV-2 infection occurs with ageing

Limited or delayed production of IFNs has been associated with a dysregulated immune response and the development of cytokine release syndrome in COVID-19^5,29^. In line with this, cytokine expression was highest in hNECs from older adults under both infected and uninfected conditions (adjusted *p*-value<0.05; **Fig. 2D**). While proteins involved in innate immune cell processes were identified in mass spectrometry-based profiling of the extracellular secretome, cytokines could not be detected (**Fig. 2E, Table S9**). Nine cytokines were detected in both mock and infected media using a 34-cytokine multiplex ELISA, with higher baseline IL-8 levels in older adults compared to young adults and children and significantly more IL-1α in secretome from older adult cultures (**Fig. 2F**). The increased baseline inflammation in older adults is aligned with the presence of inflammaging^30^. Consistent with this and with previously described alterations in ageing lungs^31^, we identified 91 unique DEGs and 372 differentially expressed proteins in uninfected hNECs between the three age groups which were enriched for canonical pathways involved in cellular metabolism, stress response, and immune and inflammatory signalling (**Fig. 2G, Table S10-13, Extended Results**).

### SARS-CoV-2 viral load is not associated with expression of SARS-CoV-2 entry factors

To identify factors that may cause differences in viral load among the age groups, we first assessed the host cell surface factors that influence viral recognition and internalisation. Ciliated and goblet cells are known to have the highest expression of SARS-CoV-2 entry receptors^12^. All cultures developed pseudostratified polarised epithelium composed of the major epithelial cell types. The virus was able to infect both ciliated (AcTub stain) and mucus secreting goblet cells (MUC5AC stain; **Fig. 3A**). Cell-type composition estimation using RNA-seq data showed an age-associated trend of higher abundance of mucus producing goblet cells and a lower abundance of ciliated cells in uninfected and infected cultures, suggesting a higher secretory phenotype with ascending age (**Fig. 3B**). The same trend was observed in the proteome with significantly lower expression of ciliated cell protein markers in older adults (adjusted *p*-value<0.005, **Fig. 3C**).

Expression of retinoic acid-inducible gene I-(RIG-I-) like receptors – crucial pattern recognition receptors which play essential roles in recognizing intracellular viral RNA – increased post SARS-CoV-2 infection in child and young adult cultures (**Fig. 3D**). With the addition of toll-like receptors (TLR), a non-significant trend of increased mean expression of these receptors were observed with ascending age (**Fig. 3D**). Recent studies suggest the SARS-CoV-2 entry factors ACE2 and TMPRSS2 ^32^ are more highly expressed on goblet cells compared to ciliated cells ^12,33^. Although ACE2 showed 20% higher expression in older adults compared to child hNECs, this difference was not statistically significant (**Fig. 3E**). Viral gene expression and *ACE2* expression were not significantly correlated, with a negative correlation between *TMPRSS2* expression and viral gene expression (Spearman’s *ρ*=-0.678, *p*<0.05; **Fig. 3F**). We then assessed other host factors potentially involved in entry of SARS-CoV-2 or those that are known to be involved in entry of other coronaviruses^12,34–36^ and observed that expression was highest in older adults, followed by children then young adults (adjusted *p*<0.005) (**Fig. 3E**). No significant difference in expression of these factors was observed between child hNECs and hBECs (**Fig. 3E**). These data suggest that physiological differences in the expression of viral recognition and entry factors are poorly correlated to SARS-CoV-2 viral load and do not explain differences in viral entry and replication associated with ageing.

### Declining transcriptional fidelity leads to subdued antiviral response to SARS-CoV-2 with ageing

As obligate intracellular parasites, viruses require host cell components to replicate, translate and transport their proteins. We assessed the expression of host proteins that interact with the SARS-CoV-2 replication and transcription complex (RTC) and may impact its transcription^37^. Using a high-confidence protein-protein interaction map between host and SARS-CoV-2 proteins^38^, we assessed the gene expression of those proteins that interact with SARS-CoV-2 RTC. Mean gene expression of host proteins that interact with the RNA-dependant RNA polymerase (Nsp12) and its binding auxiliary factor (Nsp8) was significantly higher in hNECs from children compared to those from older adults following SARS-CoV-2 infection (**Fig. 3G**). A moderately positive correlation between viral read counts and Nsp8 (Spearman’s *ρ*=0.491, *p*<0.05) and Nsp12 (Spearman’s ρ=0.418, *p<*0.05) interacting protein gene expression was observed (**Fig. 3H**). These results suggest that age-dependent differences in the expression of SARS-CoV-2 interacting proteins may influence viral transcription. SARS-CoV-2 infection, through the function of Nsp16, disrupts host mRNA splicing which aids in the suppression of the interferon response^28^. Therefore, we investigated whether susceptibility to Nsp16-mediated RNA splicing suppression occurs in an age-dependent manner. While SARS-CoV-2 infection resulted in a significant increase in the proportion of retained introns in host mRNAs in child cultures (adjusted *p*-value<0.05), there were no significant differences between age groups (**Fig. 3I**). Furthermore, retained intron proportions were moderately correlated with SARS-CoV-2 transcript expression (Spearman’s *ρ*=0.486, *p*<0.05; **Fig. 3J**), suggesting that Nsp16-mediated RNA splicing suppression occurs in a viral dose-dependent manner.

During ageing, a decline in transcriptional fidelity is thought to contribute to the senescence-associated secretory phenotype and low-grade chronic inflammation ^39^. This idea is consistent with the increased inter-cell transcriptional heterogeneity observed in the human lung with ageing ^31^. While inter-cell transcriptional heterogeneity could not be determined with bulk RNA-seq, our results showed inter-sample transcriptional heterogeneity increased with ageing under both mock and SARS-CoV-2 infected conditions (adjusted *p*-value<0.05; **Fig. 3K**). The age-related increase in transcriptional heterogeneity could result in a varied and unpredictable response to SARS-CoV-2 infection. An unsupervised principal coordinate analysis on both the transcriptome and proteome illustrated this (**Fig. 3L**). It showed that directional shifts post-SARS-CoV-2 infection were inconsistent in older adults compared to child and young adults, and Euclidean distances between centroids were smaller in older adults (0.0257) compared to child (0.0357) and young adult (0.0437). Therefore, the inherent age-related alterations in transcriptional control may lead to an inconsistent host response that is ineffective in launching an appropriate antiviral defence.

### Age-specific transcriptomic response to SARS-CoV-2 influences CoVID-19 therapeutic targets

Drug repositioning has gained considerable interest during the COVID-19 pandemic to accelerate the identification and approval of beneficial therapeutic drugs. We assessed whether the age-specific transcriptional response of hNECs to SARS-CoV-2 infection may influence the efficacy of therapeutic agents by implementing a modified version of the *COVID-19 Combinational Drug Repositioning* (COVID-CDR) platform^40^. To screen for potential repositionable drugs across different age groups, we calculated the network proximity between drug targets and SARS-CoV-2-interacting host proteins as an unbiased measure of drug-disease relatedness. Drugs (drug targets) that are more proximal (closer) to disease agents (SARS-CoV-2 interacting proteins) are theoretically more effective than more distant drugs. Drugs were considered significantly proximal to SARS-CoV-2 interacting proteins if z-score < −2.0 and *p*-value < 0.01 (**see Supplementary Information 1**).

We screened over 21,000 drugs listed in the DrugBank^41^ and identified 108 significantly proximal drugs from 39 unique drug classes across all age groups (permutation *p-*value < 0.01; **Fig. 4A-C, Table S14**). Of these drugs, 52 have evidence for treating COVID-19 symptoms or are currently in clinical trials to treat COVID-19 (**Table S14)**. Older adults possessed the greatest number of proximal drugs, with child possessing the least (**Fig. 4A**). Drug classes with more than 5 proximal drugs in all age groups were zinc compounds, antibiotics and anticancer drugs (**Fig. 4B**). Within the anticancer drug class, the majority were tyrosine kinase inhibitors which were highly proximal to several SARS-CoV-2 proteins (**Fig. 4C**). The trace metal drug classes of copper compounds and zinc compounds were identified in all age groups. In contrast, calcium compounds and aluminium compounds were exclusively identified in older adults. These results highlight that differences in the transcriptional response to SARS-CoV-2 due to ageing may impact the efficacy of therapeutic drugs for COVID-19 treatment.

**Figure 4.**
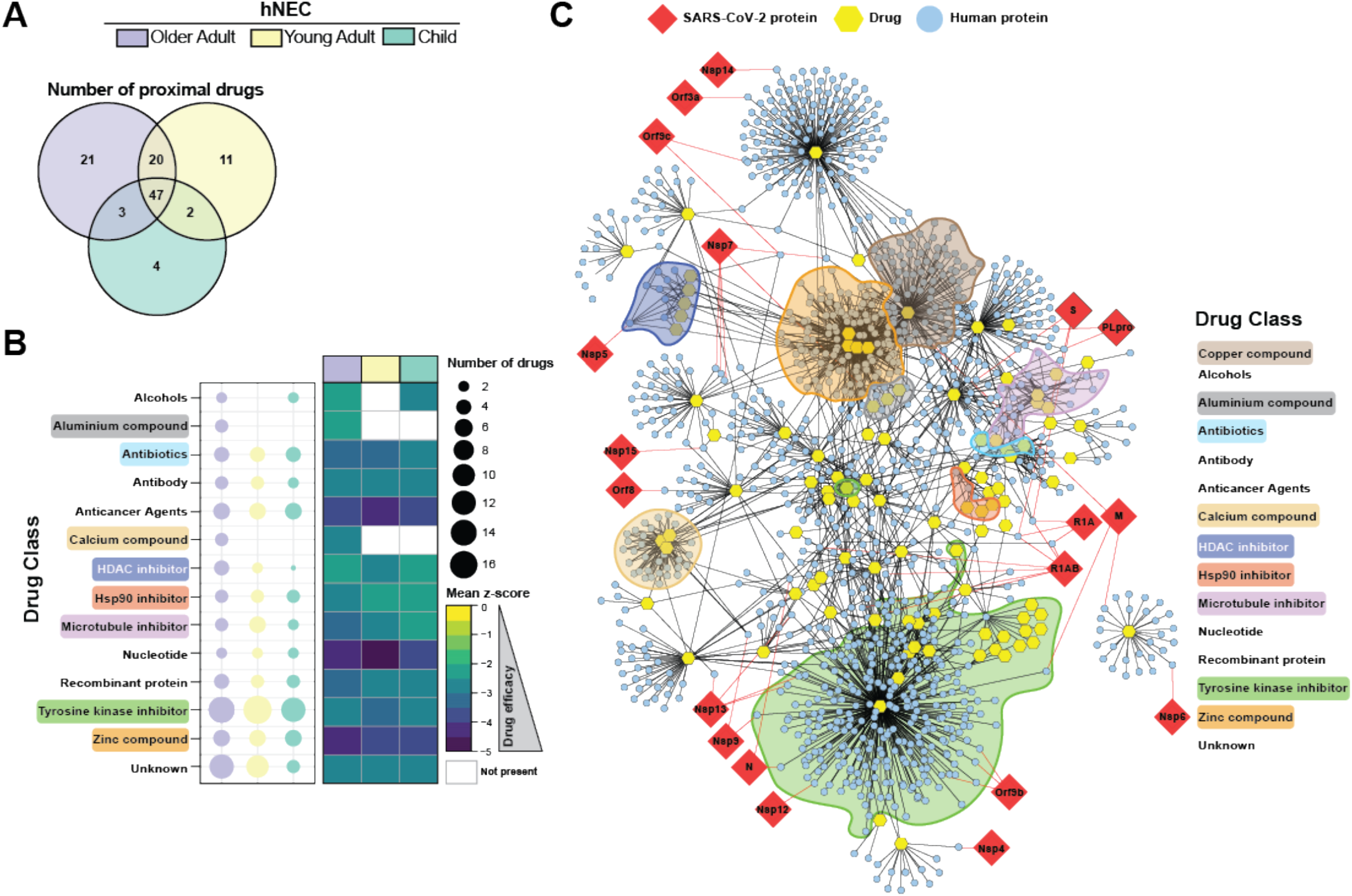
Age-specific transcriptomic response to SARS-CoV-2 influences potential drug repositioning. **(A)** Number of significantly proximal drugs (proximity *p*-value < 0.01) to SARS-CoV-2 interacting proteins in older adult, young adult and child hNECs. **(B)** Drug classes with greater than two significantly proximal drugs. Size of bubble indicates number of significantly proximal drugs in each age group. Heatmap shows the mean proximity z-score of drugs within each drug class. Lower z-scores indicates shorter proximity between drug class and SARS-CoV-2 interacting proteins. Z-score indicates drug efficacy. **(C)** Drug-target and SARS-CoV-2-human protein interaction network of all significantly proximal drugs across the older adult, young adult and child hNECs. Drug classes with drugs that were highly localised are highlighted. See **Extended Figure 4A** for network with annotated drugs and proteins. Network was produced with Cytoscape.

## Discussion

Our study revealed that viral load and antiviral responses in nasal epithelium wane with advancing age. These differences do not result from the depletion of virus-susceptible cell populations or reduced expression of the SARS-CoV-2 receptor and entry factors. Rather, the expression of host proteins hijacked by the SARS-CoV-2 replication complex are significantly reduced with ageing. The induction of innate immunity during SARS-CoV-2 infection is dependent on viral growth kinetics, viral load and MOI. With a low MOI of 0.2 the antiviral IFN response was robustly induced in child and young adults, but not in older adults. Viral load alone is unlikely to explain differences in IFN induction as bronchial epithelial cells from children exhibited a robust innate immune response despite similarly low viral load as older adult hNECs.

IFN responses are crucial for antiviral defence and in determining disease severity of COVID-19 ^26^ however SARS-CoV-2 and other coronaviruses are weak inducers of both type I and III IFN expression compared to other RNA viruses^5,26^. Relative to the nasal epithelium of children and young adults, the IFN response was severely blunted in older adults with diminished expression of ISGs. RNA processing ISGs were lowly expressed in older adults suggesting that ageing epithelial cells may be more susceptible to viral strategies that disrupt host RNA transcription and processing. In contrast to older adults, both child and young adults executed type III IFN response accompanied by upregulation of IRFs and antiviral ISGs. Young adult hNECs were the only age group with a significant upregulation of type I IFNs (IFN-β) in response to SARS-CoV-2 infection. IFN-β is regulated in a feedback loop and the upregulation of its key negative regulators (*USP18* and *SOCS3*) in young adults is likely indicative of a controlled response^44,45^. Since type I IFN receptors (IFNAR) are ubiquitously expressed, this may result in a more systemic response than that elicited by type III IFNs.

Due to the single infection timepoint we could not determine whether interferon response is delayed or impaired in older adults. However, both impaired and delayed induction of IFNs elevate inflammation in the form of excessive proinflammatory cytokine production^26^. In support, we observed significantly higher expression of proinflammatory ISGs and cytokines such as neutrophil and monocyte chemoattractive chemokines in older adult derived hNECs compared to young adult and children. This resembled the “cytokine storm” associated with severe cases of COVID-19 ^17^. High congregation of neutrophils, key cells responsible for acute lung injury in COVID-19, could explain the increased incidence of lung injury in adults infected with SARS-CoV-2. Moreover, type III IFNs, which was absent in older adults, act negatively on neutrophils, suppressing their migration and invasion^46^ and reducing the formation of neutrophil extracellular traps^47^ and the excessive production of reactive oxygen species (ROS)^48^; all of which occur in severe COVID-19^49,50^. Additionally, the decreased IFN response in older adults could cause low immunity and increased chances of reinfection. This would have further implications on the efficacy of a SARS-CoV-2 vaccine in older adults. In fact, both type I and III IFN responses stimulate antibody production and dendritic cell activity, playing an important adjuvant role for vaccine response^51^. On the other hand, young adults with both I and III IFN responses could present a better immunological memory after infection or vaccination.

We postulate that intrinsic changes in gene expression that occur with ageing alter the epithelial microenvironment in a manner that primes the cell to launch a cytotoxic proinflammatory and IFN-independent antiviral response. In support of this, we found that inter-sample transcriptional heterogeneity increased with ageing. A decline in transcriptional fidelity is involved in the development of the senescence-associated secretory phenotype that contributes to inflammaging^39^. Both transcriptomic and proteomic analysis of cultures under mock infection suggests that airway inflammaging is retained in our model. The impairment of antiviral interferon responses is thought to occur as a consequence of this mild chronic inflammatory state^52^. Furthermore, transcriptomic changes associated with ageing may impact drug targets. *In silico* analyses identified that the proximity of many therapeutic drug classes to SARS-CoV-2 interacting proteins are similar between ages, suggesting effectiveness of them to be age-independent. Yet, identification of drug classes that are exclusively present in one age group, warrant consideration for successful discovery of therapeutics and repositioning of medications for COVID-19. While a larger cohort of participants, including those who have recovered from COVID-19, will be required to confirm our findings, our data offers a foundational view of the molecular changes associated with ageing that could help yield critical diagnostic markers for managing effective therapies.

