## Supplementary Information for "Ageing impairs the airway epithelium defence response to SARS-CoV-2"

### Supplementary Information 1 – Material and Methods

#### Airway epithelial cell culture.

Human nasal airway epithelial cells (hNEC) used in this study were obtained pre-SARS-CoV-2 from brushing the nasal inferior turbinate of 6 children, 4 young adults and 3 older adults. Matched Bronchial Epithelial Cells (hBEC) were available from five children cultured from their bronchoalveolar lavage fluid (BALF). All participants and/or their carers provided written informed consent. This study was approved by the Sydney Children's Hospital Ethics Review Board (HREC/16/SCHN/120). Demographic for participants used in this study are listed in **Table S1**.

#### Conditionally reprogrammed epithelial cell (CREC) culture.

Primary basal airway cells were isolated from the participants and expanded for 1 passage on NIH 3T3 fibroblast feeders using F-medium (67.5% DMEM-F, 25% Ham's F-12, 7.5% FBS, 1.5 mM L-glutamine, 25 ng/mL hydrocortisone, 12 5 ng/mL EGF, 8.6 ng/mL cholera toxin, 24 mg/mL Adenine, 0.1% insulin, 75 U/mL pen/strep) with ROCK Inhibitor (10 μM) and antibiotics (1.25 mg/mL amphotericin B, 50 mg/mL gentamicin) as we previously described[1]. At confluence, cells were harvested by a differential trypsin method. Cells were either cryopreserved, or directly utilized for the 3D-differentiated ALI cultures. All experiments were carried out on cells passaged once.

#### 3D- Air-Liquid Interface (ALI) differentiated epithelial cell culture.

CRECs (~ 2x10^5) were seeded onto Collagen I (Advanced Biomatrix Purecol 5005) coated 6.5 mm 0.4 μM Transwell porous polyester membranes (Sigma CLS3470) in PneumaCult Ex-Plus Expansion media (Stemcell 05040). Confluent monolayers were airlifted by removing the media from the apical side and differentiated using complete PneumaCult ALI media (Stemcell 05001) in the basal chamber. Medium in the basal chamber was changed every 2 days (750 μl). HAECs were maintained at ALI for 21 days and were monitored by light microscopy for mucociliary differentiation.

#### Viral propagation and ALI infection.

The SARS-CoV-2 isolate was obtained from a nasal pharyngeal aspirate and was propagated on Vero E6 (ATCC® CRL 1586) cells in MEM (Gibco), supplemented with glutamax, penicillin (10,000 IU/mL) and streptomycin (10,000 IU/mL) and 10% FBS. Stocks were produced by infecting Vero E6 cells at a multiplicity of infection (MOI) of 0.02 and incubating the cells for 72 h. The culture supernatant was cleared by centrifugation and stored in aliquots at −80°C. Stock titers were determined using previously described methods[2]. Paired cultures for each participant were apically treated with 10 μl of PBS (Mock uninfected control) or with SARS-CoV-2 at MOI of 0.2. Unbiased inoculation with the virus was ensured by including multiple replicates in each age group. This included repeating differentiation culture for 2 of 6 child participants, 1 of 3 young adult participants and 3 of 3 older adult participants (**Table S1**). For mock samples, 1 of 3 young adult participants and 2 of 3 older adult participants were repeated to ensure consistent culturing. ALI cultures were incubated at 37°C 5% $CO_2$ for 72h. Cells were harvested for proteomics and transcriptomic analysis. The basolateral media was saved for cytokine, chemokine and secretome proteomics analysis. All samples were frozen at -80°C and once deactivated were processed simultaneously as described below. All work with infectious SARS-CoV-2 was performed in a Class II Biosafety Cabinet under BSL-3 conditions at the Kirby Institute, UNSW Sydney.

#### Immunofluorescence and imaging.

Airway ALI cultures were washed 72 h post infection (hpi) with PBS (Sigma D8662) for 5 min and fixed in 4% paraformaldehyde for 30 min at room temperature. Samples were then neutralized with 100 mM glycine in PBS for 30 min before permeabilization with 0.5% Triton-X in PBS for 30 min on ice. Fixed, permeabilized cells on transwell membranes were rinsed with PBS 3 times and then blocked using IF buffer (0.1% BSA, 0.2% Triton and 0.05% Tween 20 in PBS) with 10% normal goat serum for 1 hour at room temperature. Cells were incubated in primary antibodies against SARS Nucleocapsid, acetylated tubulin or MUC5AC (**Table S2**) for 48 h at 4°C. Cells were washed with IF buffer (3 times, 5 min each) and then incubated with a cocktail of Alexa Fluor conjugated secondary antibodies, phalloidin and DAPI (**Table S2**) for 3 h at room temperature. Cells were washed again with IF buffer (3 times, 5 min each) and the transwell membranes were excised from the inserts using a sharp scalpel (size 11). Membranes were mounted with Vectashield Plus Antifade Mounting medium (H-1900; Vector Laboratories, Burlingame, CA). For quantitative analysis of SARS N, 25x25 tilescan images of the entire membrane were acquired in the widefield mode using Zeiss Elyra PALM/SIM microscope (Carl Zeiss, Jena, Germany), 20x/0.5 objective. To visualize viral tropism, high resolution images were acquired using Leica SP8 DLS confocal microscope (Leica Microsystems, Wetzlar, Germany), 63x/1.4 oil objective. Images were then processed using ImageJ software (National Institute of Health, Bethesda, MD).

**Tilescan image analysis.**
Tilescan images were imported into Matlab (Natick, MA, USA) for SARS Nucleocapsid positive (SARS N+) cluster segmentation and extraction of cluster statistics. First, the background illumination profile was determined using 3x3 tiles at the top left corner of 25x25 tile images (area devoid of culture membrane). Each tile image was then normalised against this background value to render intensity values ⩾1. The mean intensity of regions with SARS Nucleocapsid negative cells were used to establish the cut-off threshold for presence or absence of viral infectivity, calculated to be 1.3. This threshold was applied to all tile images and a binary mask was generated for each tile image. The binary mask was further cleaned using Matlab bwmorph function to remove all SARS N+ clusters with area of 20 pixels or less (pixel size 0.5 μm). The percentage covered by SARS N+ was then extracted. The binary mask was also decomposed using bwlabel function into separate clusters, and the number of SARS N+ clusters, mean area as well as the mean intensity of individual clusters were extracted.

**Ciliary Beat Frequency (CBF) measurement.**
Ciliary beating frequency was measured for each participant's mature ALI cultures prior to infection experiments. Cilia beating were imaged in transmission light modality using ORCA-Flash 4.0 sCMOS camera (Hamamatsu Photonics, Shizuoka Pref., Japan), connected to Zeiss Axio Observer Z.1 inverted microscope (Carl Zeiss, Jena, Germany). Images were acquired serially at 334 frames per second (fps) with 3 ms exposure time on a EC Plan-Neofluar 20x/0.5 Ph2 M27 dry objective (512 x 512 pixel; 0.325 μm x 0.325 μm). Six time-series images were sampled at random from triplicate filters per treatment condition. Imaging was performed after stabilisation of filters for 10 min at 37°C, 5% $CO_2$ and under this maintained physiological environment. To extract cilia beating spectra and corresponding beating frequency, image series were analysed using a custom-built script in Matlab (MathWorks, Natick, MA) as described previously[1]. The dominant frequency (a frequency with the highest peak) within the physiological range of cilia beating (3-30Hz) was identified using the Matlab function 'findpeaks'. A paired two-tailed Student's t-test was performed to determine statistical significance between age groups ($p < 0.05$) using Prism 8 (v8.4.1).

**Real-Time Quantitative PCR of SARS-CoV-2.**
Real-Time Quantitative PCR of SARS-CoV-2. Isolated RNA from mock and SARS-CoV-2 infected ALI cell lysate (1 ng) and basolateral media (10 ng) was used as input for the AllPlex SARS-CoV-2 multiplex real-time PCR assay (Seegene CAT RV10248X) to detect three SARS-CoV-2 genes: E gene, rdRP/S gene and N gene. C(t) values were normalised to HEX fluorophore internal control. The virus copy numbers per reaction were quantified using a control plasmid which contained the complete nucleocapsid and envelope gene from SARS-CoV-2. Unpaired two-tailed Student's t-test was performed to determine statistical significance between age groups ($p < 0.05$) using Prism 8 (v8.4.1).

**Harvesting of cells for bulk RNA-Sequencing.**
In total, 46 mature mock and infected ALI cultures were harvested for bulk RNA-sequencing. Samples were homogenized with 1 mL of cold TRIzol (Life Technologies) for RNA extraction. RNA purification was performed with a RNeasy Mini Kit (Qiagen) following the manufacturer's protocol

with an on-column RNase-free DNase Set (Qiagen) treatment. RNA quality was analyzed using the 2100 Bioanalyzer (Agilent Technologies). For each sample, 300 ng of high-quality RNA (RIN > 9) was polyA-selected and sequencing libraries prepared using the TruSeq RNA Library Prep Kit v2 (Illumina) or TruSeq Stranded mRNA Library Prep Kit (Illumina) according to the manufacturer's instructions. Sequencing libraries were sequenced on an Illumina NextSeq 500 platform with 76 bp single-end reads at the Ramaciotti Centre for Genomics (UNSW Sydney, Australia). Expression values of genes are available at Github repository (https://github.com/avatar-1/age-cov) and raw sequence data are available upon request after proper approval processes.

**Quality control and alignment of reads.**

Data quality metrics were calculated using FastQC (v0.11.9)[3] and summary statistics plots were visually inspected. The human reference genome containing decoy sequences but lacking alternate contigs (GRCh38, GCA_000001405.15_GRCh38_no_alt_plus_hs38d1_analysis_set.fna) was merged with the SARS-CoV-2 genome (NC_045512.2) for alignments. The human Y chromosome derived sequences were hard masked in the reference genome for alignments of female samples to ensure accuracy for RNA quantitation[4]. Subread (v2.0.1)[5] was used for alignments in RNA-seq mode (parameters: -t 0) and multimapping reads were assigned to one random location (parameters: --multiMapping -B 1). RefSeq annotations for the human genome downloaded from the UCSC database (hg38.ncbiRefSeq.gtf.gz) and the SARS-CoV-2 genome annotations downloaded from the RefSeq database (GCF_009858895.2_ASM985889v3_genomic.gtf.gz) were merged and used during alignment (-a hg38.sarscov2.refseq.gtf). Subsequently, featureCounts within the Subread package was used to count reads that overlapped genomic features (-O -M -s 2) to ensure that multimapping reads are counted at least once against a feature. Reads were assigned to all of their overlapping features.

**Differential gene expression analysis.**

Raw counts were normalized using edgeR (v3.30.3)[6] using the trimmed mean of M-values (TMM) method [7]. Genes with expression values in top 99.5% in any one sample were retained for the analyses. Differential gene expression analysis was performed with edgeR in a pairwise manner between treatment conditions using the generalized log-linear model (glm) method. For differential expression analysis between age-groups within conditions, each age-group was compared to the combination of the other two age-groups. *P*-values were adjusted using the Benjamini-Hochberg procedure to calculate false discovery rates (FDR). Genes with log2 fold-change greater than |0.585| (fold-change > |1.5|) and FDR < 0.05 were considered significantly differentially expressed. All graphs were plotted in R (v4.0.0)[8] using RStudio (v1.2.5033) and ggplot2 (v3.3.2)[9]. Differential expression analyses were automated using an RScript available at https://github.com/avatar-1/age-cov.

**Co-expression analysis.**

Pearson correlation was performed to measure the co-expression of up/down-regulated long non-coding RNAs (lncRNAs) with mRNAs (correlation coefficient > 0.8). The putative function of each lncRNA was estimated based on biological functions enriched by the co-expressed mRNAs as per the standard guilt-by-association assumption. Functional enrichment of co-expressed mRNAs was performed using the right-tailed Fisher's exact test (hypergeometric null distribution) followed by FDR correction (FDR < 0.05). The analysis was performed in R using an in-house script available at https://github.com/avatar-1/age-cov.

**Cell lineage analyses.**

Estimation of the abundance of epithelial cell-types within bulk RNA-seq data was performed with CIBERSORTx (accessed 28 August 2020) [10]. Normalized single-cell RNA sequencing gene expression counts of ALI-differentiated human bronchial epithelial organoids (GSE102580)[11] were used as a single cell reference sample using default settings. Cell fraction imputation was performed with quantile normalization disabled and 1000 permutations for significance analysis. Unpaired student's t-test with Bonferroni correction was used to determine statistical significance (adjusted *p*-value < 0.05) between age-groups and condition using the rstatix (v0.6.0) package in R[12]. Graphs were plotted in R using ggplot2.

**Lung age-associated gene comparison.**
Gene expression of a database of high confidence age-associated genes in the human lung from Chao *et al.* [13] was compared between age groups. Only genes with expression of greater than zero raw reads across all samples were retained. Only proteins present in over 50% of replicates were included. A two-tailed Wilcoxon rank sum test with Bonferroni correction was used to determine statistical significance (adjusted $p$-value < 0.05) between age groups using the rstatix package in R. Graphs were plotted in R using ggplot2.

**Interferon stimulated genes and cytokine comparative analysis.**
Gene expression of a database of high confidence interferon stimulated genes determined by Mostafavi *et al.*[14] and a list of cytokines (chemokines and interleukins) was compared between age groups. Only genes with expression of greater than zero raw reads across all samples were retained. A two-tailed Wilcoxon rank sum test with Bonferroni correction was used to determine statistical significance (adjusted $p$-value < 0.05) between age groups using the rstatix package in R. Graphs were plotted in R using ggplot2.

**Transcriptional noise analysis.**
Transcriptional noise was calculated as previously described for single-cell RNA sequencing experiments [15,16]. Briefly, genes with greater than 0 raw reads across all samples were retained and were binned into 10 bins based on the mean expression, with the top and bottom bins excluded. The coefficient of variation (defined as the standard deviation of expression divided by the mean expression) was calculated and 10% of the genes with the lowest coefficient of variation retained. Retained raw reads were log2 transformed. The Euclidean distance between each sample and the sample mean within each age group was calculated and used as a measure of transcriptional noise. A two-tailed Wilcoxon rank sum test was used to determine statistical significance ($p$<0.05) between groups using the rstatix package in R. Graphs were plotted in R using ggplot2.

**Principal coordinate analysis.**
Principal coordinate analysis (PCoA) was performed with the plotMDS function in EdgeR. Permutational multivariate analysis of variation was performed using the *vegan* (v2.5-7) R package[17] with the adonis function with 999 permutations. Multivariate homogeneity of group dispersion was calculated using the *vegan* R package with the vegdist and betadisper functions. Distances between the groups were performed with the distance between centroids function within the *usedist* (v0.4.0) R package [18]. Graphs were plotted in R using ggplot2.

**Intron retention analysis.**
Raw sequencing reads were aligned to the human reference genome (GRCh38, GCF_000001405.39_GRCh38.p13_genomic.fna) using STAR (v2.7.2b)[19] with default settings. Alignment files (.bam) outputted by STAR was used as inputs for the alternative transcript splicing analysis tool rMATS (v4.0.2)[20] which was run with default settings. Raw retained introns (RI) counts identified by rMATS outputted in the JC.raw.input.RI.txt file was used for further analysis. Specifically, the number junction reads that span the intron-exon boundary (IJC) and the number of junctions reads that skip the exon-intron boundary (i.e., span the exon-exon boundary; SJC) were retained. Junction reads were normalised against the total read count of each sample. RI proportion was calculated by: *IJC / (IJC + SJC)*. RI proportion was calculated for each RI event identified by rMATS. The RI proportion across all host genes was plotted for all samples as violin plots in R using ggplot2. A two-tailed Wilcoxon rank sum test was used to determine statistical significance (adjusted $p$-value < 0.05) between groups using the rstatix package in R.

**Virus-host-drug multi-dimensional network construction.**
To reveal the interplay between COVID-19 drug targets, and SARS-CoV-2-human protein interactions on human protein-protein interactome, a multi-dimensional network was fused from different interactome data sources integrating drug-target, SARS-CoV-2-human, and human protein-protein interactions (PPIs) into a single network. PPIs in humans were downloaded from 'Interologous Interaction Database' (I2D), version 2.9[21]. High-confidence protein–protein interactions between

SARS-CoV-2 and human proteins were obtained from Gordon *et al.* [22] and Saha *et al.* [23]. Drug-target interactions were compiled from DrugBank, version 5.1.6. Interactions between SARS-CoV-2 and human proteins were collected by searching through literature (see COVID-CDR[24] for more details). Genes expressed across each age group from the hNEC-SARS-CoV-2 infected transcriptome data, were selected as those whose average expression across replicates greater than or equal to the mean expression (**Extended Figure 4B**). SARS-CoV-2-host-drug network was then constructed by filtering out host proteins that are not expressed within the age group. The corresponding filtered networks were used for subsequent drug-SARS-CoV-2 network proximity estimation.

**Network-based topological proximity measure.**

As previously proposed [25], to capture the network proximity between drug (A) and disease (C), we used the average shortest path length between disease proteins to the nearest target of drug (A) on human PPI network using Equation below, where $T = \{t\}$ is the set of targets of a drug, $C = \{c\}$ is the set of COVID-19-related proteins, and $d(t,c)$ is the shortest distance between a target $a$ and a disease protein $c$.

$$d_{TC} = \frac{1}{\|T\| + \|C\|}\left(\sum_{t\in T}\min_{c\in C} d(t,c) + \sum_{c\in C}\min_{t\in T} d(t,c)\right)$$

The proximity measures were then converted to z-scores (i.e., $z = \frac{d_{TC} - \mu}{\sigma}$) by comparing the observed distance to a reference distance distribution ($\mu$ and $\sigma$) obtained by permutation test of 100 iterations where at each iteration a randomly selected group of proteins of matching size was generated as the disease proteins and drug targets in the human interactome. The corresponding $p$-value was used to screen for drugs with significant proximity ($p$-value < 0.01) to SARS-COV-2 related proteins on human PPI network. Drug-target and SARS-CoV-2-human protein interaction network was created in Cytoscape[26].

**Cell lysate preparation for label-free mass spectrometry proteomics.**

For the mass spectrometry analysis, total protein from one transwell per participant was extracted by homogenising the cells in 100µl of RIPA buffer (Life Technologies 89900) containing protease inhibitor cocktail (Sigma 11836153001). Samples were sonicated using the Bioruptor Pico (Diagenode B01060010) for a total of 20 cycles of 30 sec on/30sec off at 4°C. Protein concentrations were determined using the 2-D Quant kit (GE Life Sciences 80648356). Samples were reduced (5 mM DTT, 37C, 30 min), alkylated (10 mM IA, RT, 30 min) then incubated with trypsin at a protease:protein ratio of 1:20 (w/w) at 37°C for 18 h. The peptide solution was desalted with two SDB-RPS disk (Empore, Sigma Cat # 66886-U) packed in the 200 µl pipette tip following the previous published procedure[27] . Eluted peptides from each clean-up were evaporated to dryness in a SpeedVac and reconstituted in 10 µL 0.1% (v/v) formic acid, 0.05% (v/v) heptafluorobutyric acid in water.

**Basolateral medium preparation for label-free mass spectrometry of secretome.**

Basolateral media (200 µl) collected at 72 hpi from control and SARS-CoV-2 airway ALI cultures were prepared for analysis by NanoLC MS ESI MS/MS (LC-MS/MS). The virus was deactivated with an equal volume of RIPA buffer (Life Technologies 89900) containing protease inhibitor cocktail (Sigma 11836153001). Samples were sonicated using a Bioruptor Pico (Diagenode B01060010) for 10 cycles of 30s on/30 off at 4°C. Soluble proteins were precipitated by addition of ten volumes of 2% (w/v) trichloroacetic acid (TCA) in isopropanol and vigorously shaking for 5 min at RT. Samples were centrifuged at 14,000 x g for 10 min at 4°C to pellet proteins. Pellets were washed twice with methanol and allowed to air dry. Protein pellets were then re-dissolved in RIPA buffer (Thermo Fisher), sonicated for 5 min in a water bath followed by heating at 80°C for 10 min before reduction (5mM DTT at 37°C for 30 min) and alkylation (10mM iodoacetamide for RT for 30 min in the dark). Sequencing grade trypsin (Promega) 0.2 µg was added and incubated overnight at 37°C. The peptide solution was desalted with the SDB-RPS procedure described above. Eluted peptides from each clean-up were evaporated to dryness in a SpeedVac and reconstituted in 10 µL 0.1% (v/v) formic acid, 0.05% (v/v) heptafluorobutyric acid in water.

**Mass Spectrometry.**
Proteolytic peptide samples were separated by nanoLC using an Ultimate nanoRSLC UPLC and autosampler system (Dionex, Amsterdam, Netherlands) and analyzed on an Orbitrap Fusion Lumos Tribrid mass spectrometer (Thermo Scientific, Bremen, Germany) as previously described[28]. A 90 min gradient and a 60 min gradient were used for lysate and supernatant samples, respectively.

**Sequence database searches, protein identification and quantification.**
Raw peak lists derived from the cell lysate experiments were analysed using MaxQuant (v.1.6.2.10)[29] with the Andromeda algorithm[30]. Search parameters were: ±4.5 ppm tolerance for precursor ions and ±0.5 Da for peptide fragments; carbamidomethyl (C) as a fixed modification; oxidation (M) and N-terminal protein acetylation as variable modifications; and enzyme specificity as trypsin with two missed cleavages possible. Peaks were searched against the human Swiss-Prot database (August 2018 release) and SARS-CoV-2 Swiss-Prot database (August 2020). Label-free protein quantification was performed using the MaxLFQ algorithm with default parameters. Differential protein expression analysis was performed with the R package DEP (v.1.10.0)[31] within R using RStudio. Functional analysis of differentially abundant proteins was performed with IPA. Graphs were plotted in R using ggplot2. The full list of identified proteins and label-free quantification (LFQ) intensity outputted from MaxQuant is available at https://github.com/avatar-1/age-cov. Mass spectrometry data are available at the ProteomeXchange Consortium via the PRIDE partner repository with the dataset identifier PXD025059.

**Functional enrichment analysis.**
Functional enrichment analysis of differentially expressed genes and proteins was performed with Core Analysis in Ingenuity Pathway Analysis (IPA; v20.0; Qiagen, https://www.qiagenbioinformatics.com/products/ingenuity- pathway-analysis). Gene ontology enrichment analysis of gene lists was performed with gProfiler[32] (accessed August 2020).

**Multiplex cytokine and chemokine assay.**
The basolateral medium was collected at 72 hpi and assay standards were inactivated with 0.05% beta propiolactone (BPL) (Sigma-Aldrich) overnight at 4ºC. The BPL was hydrolyzed at 37ºC for 1 h prior to the multiplex assay[33]. The abundance of cytokines was measured using 34-Plex Human ProcartaPlex™ Panel 1A according to the manufacturer's instructions (ThermoFisher Scientific, Sydney, NSW, Australia). Luminex MAGPIX instrument (Luminex Corporation, Northbrook, IL, USA) calibrated with MAGPIX Calibration and Performance Verification Kits (Millipore) with xPONENT software (Luminex) was used to acquire data and analysed with Multiplex Analyst software version 5.1 (Luminex) as the Median Fluorescence Intensity (MFI). Statistical analysis was performed with an unpaired two-tailed Student's t-test using Prism 8 (GraphPad Software, San Diego, CA). $p < 0.05$ was considered statistically significant.

**Statistical analysis.**
The statistical analyses used for each experiment are stated in the respective sections.

**Code availability**

All code used to perform analyses on the transcriptomic and proteomic data is available via GitHub at https://github.com/avatar-1/age-cov.

### Supplementary Information 2 – Extended Figures

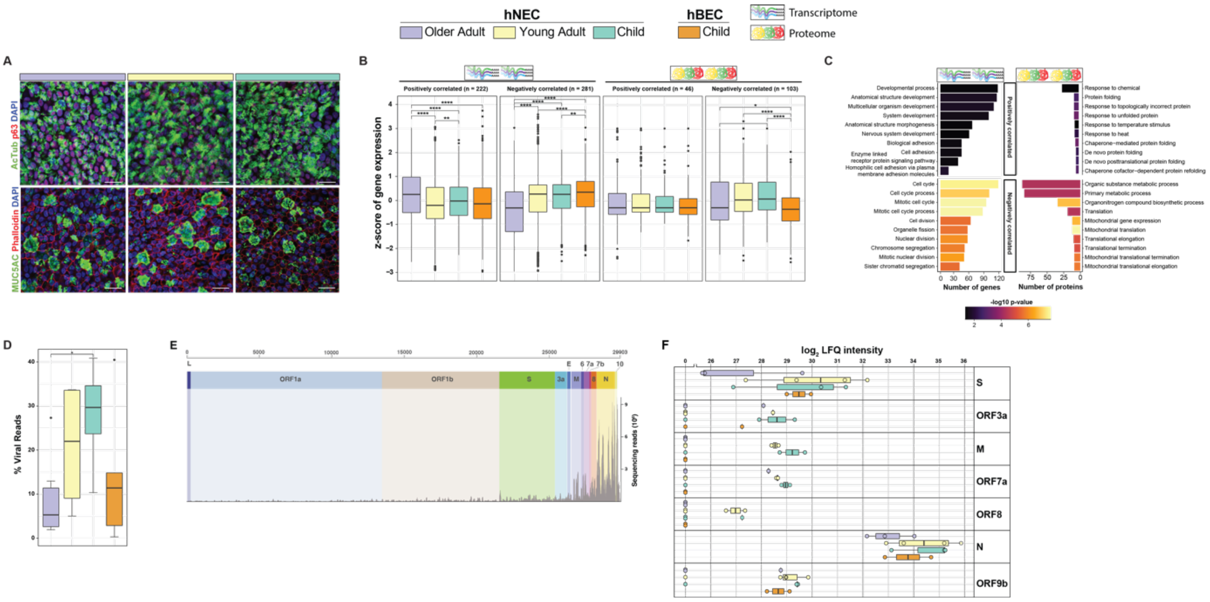


**Extended Figure 1. (A)** Representative immunofluorescence images of differentiated mock-infected hNECs stained for p63 (basal progenitor cells, red), acetylated tubulin (ciliated cells, green) (top). MUC5AC (mucus-producing goblet cells, green), phalloidin (actin, red) (bottom). DAPI (nucleus, blue). 63x/1.4 oil immersion objective. Scale bars: 20μm. **(B)** Tukey boxplot of relative gene and protein expression (z-score) of age-associated genes in the human lung identified by Chao et al. (2021) (9) in mock fully differentiated mature HAEs. Left panel shows genes positively correlated with increasing age and the right panel shows genes negatively correlated with increasing age. Mock, child hNEC, *n* = 5; child hBEC, *n* = 5, young adult, *n* = 4; older adult, *n* = 5; virus, child hNEC, *n* = 8; child hBEC, *n* = 5, young adult, *n* = 4; older adult, *n* = 6; two-tailed Wilcoxon rank sum test with Bonferroni correction. **(C)** Canonical pathway analysis of differentially expressed proteins between mock-infected hNEC age groups using IPA. Color indicates the -log10 p-value of each canonical pathway. **(D)** Raw sequencing reads aligned to the SARS-CoV-2 genome across all hNEC samples. Mock, child hNEC, *n* = 5; child hBEC, *n* = 5, young adult, *n* = 4; older adult, *n* = 5; virus, child hNEC, *n* = 8; child hBEC, *n* = 5, young adult, *n* = 4; older adult, *n* = 6; two-tailed Wilcoxon rank sum test with Bonferroni correction. **(E)** Percent of reads that aligned to the SARS-CoV-2 genome across all cultures. (**F)** Tukey boxplot of the abundance of SARS-CoV-2 proteins measured using label-free quantitative proteomics. Higher $\log_2$ LFQ intensities indicate higher abundance of a protein. Mock, child hNEC, *n* = 2; child hBEC, *n* = 2, young adult, *n* = 4; older adult, *n* = 3; virus, child hNEC, *n* = 3; child hBEC, *n* = 3, young adult, *n* = 4; older adult, *n* = 4; two-tailed Wilcoxon rank sum test with Bonferroni correction. * adjusted *p*-value < 0.05, ** < 0.01, *** < 0.001. Boxes indicate the first and third quartiles with middle horizontal line indicating the median and external horizontal lines indicating minimum and maximum values.

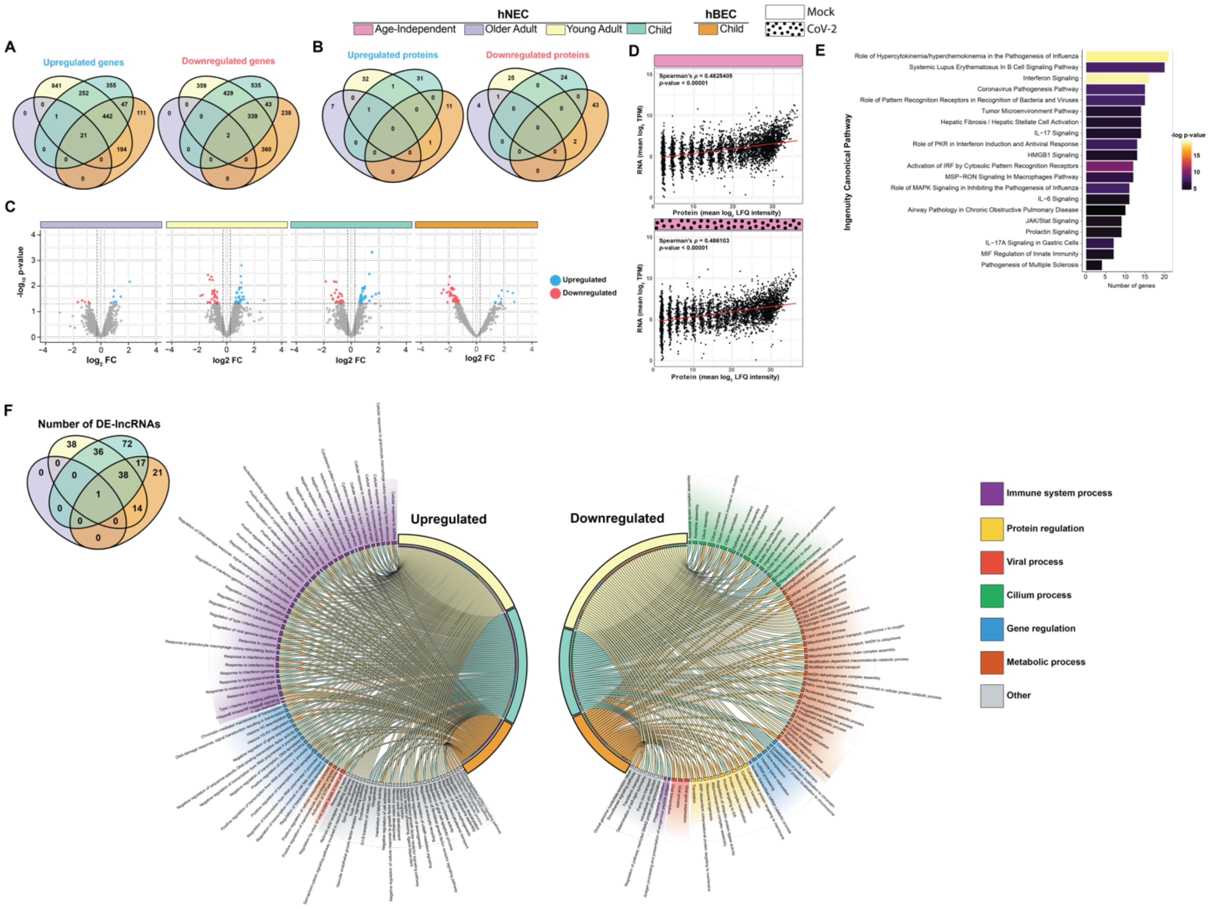


**Extended Figure 2.** Number of **(A)** genes and **(B)** proteins that are differentially expressed in common between the three age groups. **(C)** Volcano plots of differentially expressed proteins between mock and viral-infected cultures within each age group comparison. (**D)** Spearman's correlation of RNA ($log_2$ transcripts per million (TPM)) and protein (log2 label-free quantification (LFQ) intensity) expression in all mock-infected and viral-infected hNECs. (**E**) Canonical pathway analysis of less stringent cut-off (*p*-value < 0.05) differentially expressed genes in older adults by IPA. Color indicates the -log10 p-value for each pathway. * adjusted *p*-value < 0.05, ** adjusted *p*-value < 0.01, *** adjusted *p*-value < 0.001. (**F**) Venn diagram of the number of differentially expressed (DE) lncRNAs and circos plots of the enriched gene ontologies of differentially expressed lncRNAs co-expressed mRNAs within each age group.

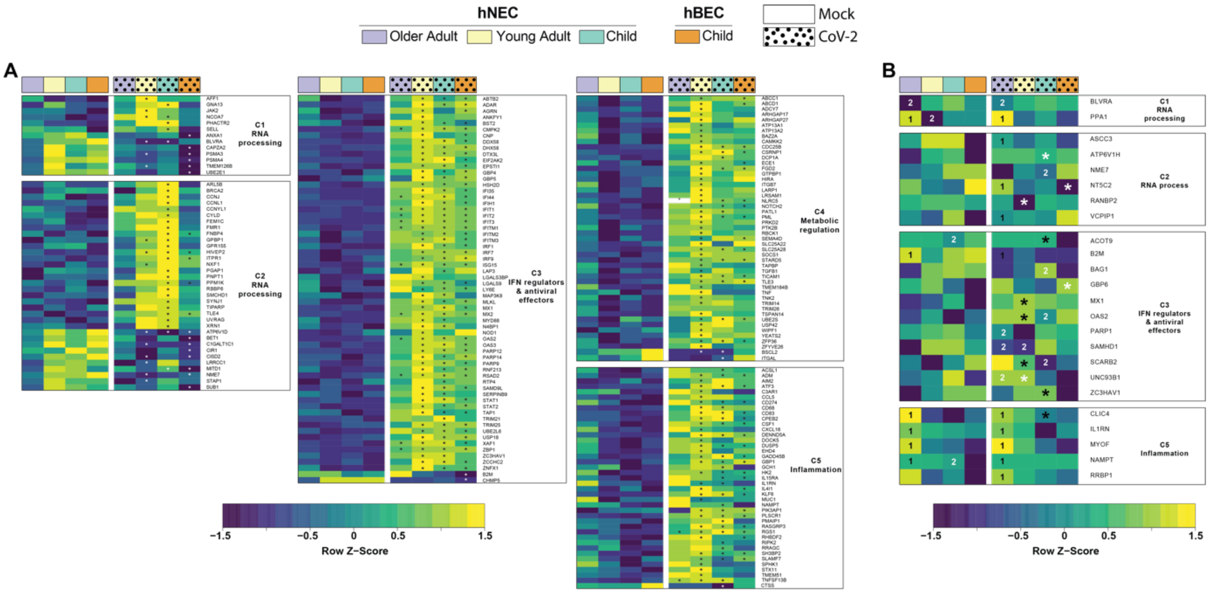


**Extended Figure 3.** Heatmap of the relative average **(A)** gene and **(B)** protein expression of interferon-stimulated genes (ISG) within functional clusters as defined by Mostafavi *et al*. Each tile in the heatmap shows the normalized expression of a gene (Z-score). Color key shows the row Z-score (yellow, high; blue, low). * significantly upregulated (black) and downregulated (white) in SARS-CoV-2 infection relative to mock. (1) significantly upregulated and (2) downregulated relative to the two other age groups within the same condition.

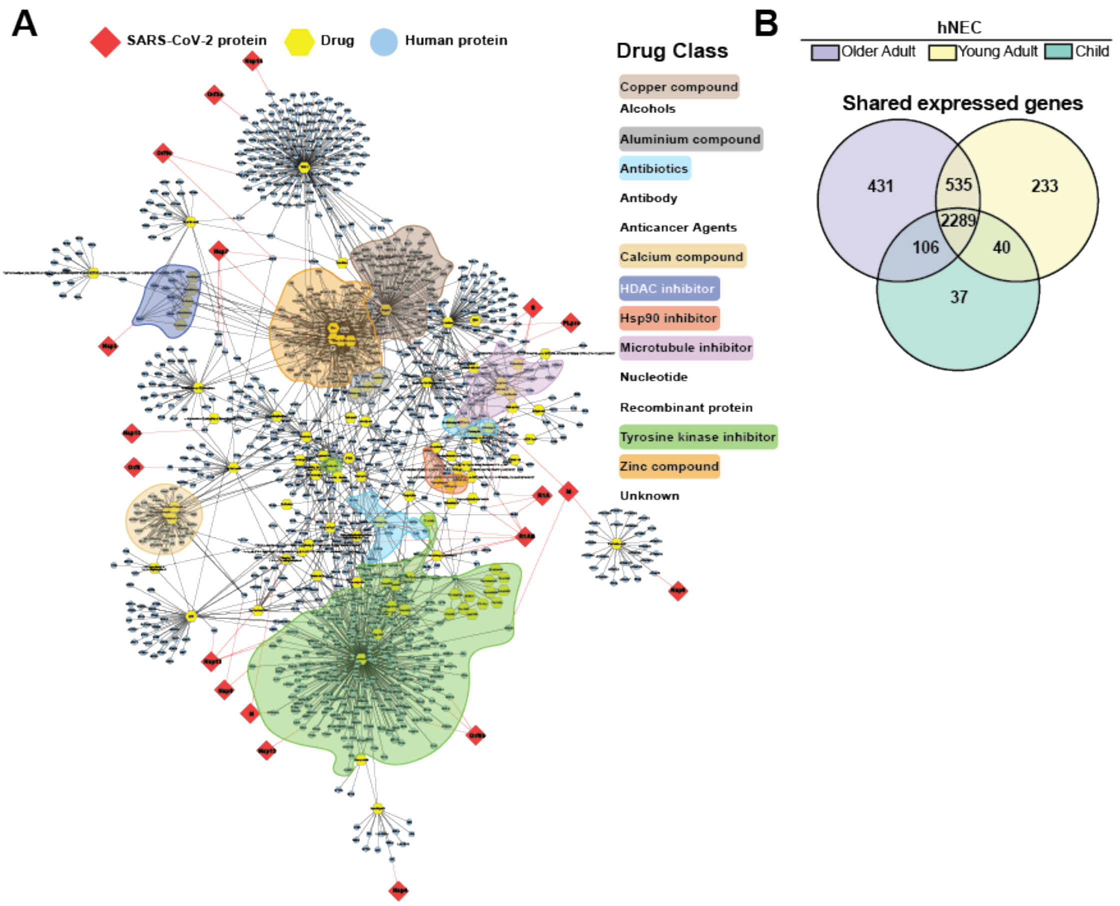


**Extended Figure 4. (A)** Drug-target and SARS-CoV-2-human protein interaction network of all significantly proximal drugs across the older adult, young adult and child hNECs. Drug classes with drugs that were highly localised are highlighted. **(B)** Venn diagram of expressed genes shared between the three age groups in hNECs. Expressed genes were defined as those with higher expression (counts per million) than the mean expression in each age group.

### Supplementary Information 3 – Extended Results

#### 1. Decreasing viral load in differentiated nasal epithelial cultures from older adult

Similar to previous reports [34,35], SARS-CoV-2 viral RNA and protein were only detected at trace levels in the basolateral medium (**Table S9**). Transcript levels of envelope (*E*) RNA-dependent RNA polymerase (*RdRp*), spike (*S*), and nucleocapsid (*N*) were significantly higher ($p < 0.01$) in hNEC lysate of child and young adult than older adults (**Fig. 1D**). RNA-seq analysis of infected hNECs showed 38% of viral RNA reads aligned to the SARS-CoV-2 structural protein-coding genes *S, E*, *M* and *N* which is indicative of full transcription and replication of the virus (**Extended Fig 1E**). Viral RNA reads ranged from 0.3% to 47% of total RNA reads (**Fig. 1E**). Significant and reproducible higher read counts of SARS-CoV-2 transcripts were observed in children compared to older adult hNEC (mean: 28% and 9%, respectively), confirming the qPCR results (**Fig. 1D, Extended Fig 1D**). In the cell lysate, out of the 26 reported SARS-CoV-2 proteins (NCBI Reference Sequence: NC_045512.2), seven (S, M, N, ORF3a, ORF7a, ORF8 and ORF9b) were detected in the proteome of viral infected cultures (**Extended Fig 1F**). Abundance of viral proteins decreased with ascending age although statistical significance was not achieved (**Extended Fig 1F**). Expectedly, qPCR detection of SARS-CoV-2 transcript in infected child hBEC was significantly lower ($p < 0.05$) than their hNEC counterparts (**Fig. 1D**). SARS-CoV-2 viral transcript expression in child HBEs was similar to that of older adult hNECs (mean: 14% and 9%, respectively). Consistent with this data, lower viral proteins were detected (**Extended Fig 1F**) and IF staining of the SARS-CoV-2 N in the infected hBECs compared to hNECs (**Fig. 1C)**.

#### 2. SARS-CoV-2 infection causes differential expression of long non-coding RNAs in child and young adults

A large number of long non-coding RNAs (lncRNA) were differentially expressed after SARS-CoV-2 infection in young adult (n = 127), and child (n = 164) hNECs and hBECs (n = 92) (**Table S8**). In contrast, only *RN7SL1* (RNA component of signal recognition particle) was differentially expressed in older adults (**Extended Fig 2F)**. Common to young adults and both child nasal and bronchial cultures, upregulated lncRNAs were co-expressed with mRNAs that were highly enriched for immune system processes including interferon signaling and antiviral response, and gene regulatory processes, while downregulated lncRNAs were co-expressed with mRNAs that were enriched for cilium and metabolic processes (**Extended Fig 2F**).

#### 3. Dysregulated IFN signalling and antiviral responses to SARS-CoV-2 infection during ageing

Seventy-two hpi with SARS-CoV-2, expression of type I (*IFNB1*) and type III IFNs (*IFNL1*, *IFNL2*) did not significantly increase in older adults (**Fig. 2A**). Type III interferons were upregulated in all other cultures, whereas upregulation of type I was only detected in young adult hNEC and child hBEC (**Fig. 2A**). Expression of the IFN regulatory factors (IRF), which regulate IFN expression, were also not significantly altered in older adult groups post SARS-CoV-2 infection. *IRF1*, *IRF7*, and *IRF9* were significantly upregulated in children (hNEC and hBEC). In young adult hNECs in addition to above, *IRF5* was also significantly upregulated (**Fig. 2A**). The type III IFN receptor gene *IFNLR1* was significantly upregulated in young adults but no other age groups (**Fig. 2A**). Neither type I nor II IFN receptor genes were differentially expressed post-infection in any cultures (**Table S4**). IFNs induce the expression of several hundred IFN-stimulated genes (ISG) that are the first-line of the antiviral defence in mammals. To understand the biological functions associated with ISGs, we annotated ISGs and grouped them based on five functional clusters as reported by Mostafavi *et al.* [14]; RNA processing (C1 and C2), interferon regulation and antiviral function (C3), metabolic regulation (C4) and inflammation

(C5; **Fig. 2C, Extended Fig 3A-B**). In mock cultures, mean expression of genes in C3, C4 and C5 clusters were significantly higher in older adults compared to young adults and children (adjusted *p*-value < 0.0001). After SARS-CoV-2 infection, mean expression of genes in C1/C2 clusters significantly decreased in child hBECs, whereas their mean expression significantly increased in hNECs of all ages, with child hNECs showing significantly higher mean expression than both young and older adult hNECs (adjusted *p*-value < 0.0001). Mean expression of all other ISG clusters significantly increased in all groups compared to their mock counterparts (adjusted *p*-value < 0.05). While mean expression of genes in clusters C3 and C4 did not differ between age groups, the mean expression of genes in the C5 cluster (inflammatory) was significantly higher in older adults than both child cultures (adjusted *p*-value < 0.0001). In older adults, this response was elicited by a very small (n = 14) subset of upregulated ISGs (**Extended Fig 3A**). In comparison, drastically higher numbers of ISGs were modulated in young adults (156), child hNEC (132) and hBECs (109). Key antiviral ISGs significantly upregulated in all age-groups included *IFIT1*, *IFIT2*, *IFIT3*, *MX2*, and *OAS2* (**Extended Fig 3A**). As only 117 ISGs could be identified in the proteome in over 50% of samples, significant differences in the mean expression of proteins within ISGs clusters could not be determined between age groups. Across all age groups, 11 differentially expressed ISGs were identified between mock and SARS-CoV-2 cultures, including MX1 and OAS2 in young adults (**Extended Fig 3B**). All proteins identified that belonged to the C5-inflammation cluster were expressed significantly higher in older adults than other age groups.

#### 4. Inflammaging and inflammatory responses to SARS-CoV-2 infection during ageing

Mean expression of cytokines were significantly higher in mock older adults compared to mock child hNECs and hBECs (adjusted *p*-value < 0.05; **Fig. 2D**). Seventy-two hpi with SARS-CoV-2, mean expression of cytokines significantly increased in all cultures with greatest significance in child hNECs. Despite significantly lower IFN expression in older adults compared to other age groups, older adults had higher mean expression of cytokines to other hNEC age groups and was significantly higher compared to child hBECs (adjusted *p*-value < 0.05; **Fig. 2D**). Chemokines of the CXC subgroup (CXCL) and interleukins (ILs) were highly upregulated in children and young adults while CC subtype (CCL) chemokines were downregulated in children (**Fig. 2D**). In older adults, only *CXCL10* and *CXCL11* were upregulated post-infection. *CXCL3*, *CXCL5* and *CXCL6*, which encode chemoattractants of neutrophils, *IL1A* and *IL1B*, which encodes acute phase pro-inflammatory cytokine, and *IL33*, which encodes a pro-T-helper-2 (Th2) cytokine, were expressed significantly lower in SARS-CoV-2 infected child hNECs than in adults (both young and older) (**Fig. 2D**). The higher expression of chemokines and ILs in young and older adults compared to children may indicate an exaggerated inflammatory response to SARS-CoV-2 infection in adults.
